# Transcriptome-guided parsimonious flux analysis improves predictions with metabolic networks in complex environments

**DOI:** 10.1101/637124

**Authors:** Matthew L. Jenior, Thomas J. Moutinho, Bonnie V. Dougherty, Jason A. Papin

**Affiliations:** Department of Biomedical Engineering, University of Virginia, Charlottesville, VA, USA; Department of Medicine, Division of Infectious Diseases & International Health, University of Virginia, Charlottesville, VA, USA; Department of Biochemistry & Molecular Genetics, University of Virginia, Charlottesville, VA, USA

## Abstract

The metabolic responses of bacteria to dynamic extracellular conditions drives not only the behavior of single species, but also entire communities of microbes. Over the last decade, genome-scale metabolic network reconstructions have assisted in our appreciation of important metabolic determinants of bacterial physiology. These network models have been a powerful force in understanding the metabolic capacity that species may utilize in order to succeed in an environment. Increasingly, an understanding of context-specific metabolism is critical for elucidating metabolic drivers of larger phenotypes and disease. However, previous approaches to use network models in concert with omics data to better characterize experimental systems have met challenges due to assumptions necessary by the various integration platforms or due to large input data requirements. With these challenges in mind, we developed RIPTiDe (Reaction Inclusion by Parsimony and Transcript Distribution) which uses both transcriptomic abundances and parsimony of overall flux to identify the most cost-effective usage of metabolism that also best reflects the cell’s investments into transcription. Additionally, in biological samples where it is difficult to quantify specific growth conditions, it becomes critical to develop methods that require lower amounts of user intervention in order to generate accurate metabolic predictions. Utilizing a metabolic network reconstruction for the model organism *Escherichia coli* str. K-12 substr. MG1655 (iJO1366), we found that RIPTiDe correctly identifies context-specific metabolic pathway activity without supervision or knowledge of specific media conditions. We also assessed the application of RIPTiDe to *in vivo* metatranscriptomic data where *E. coli* was present at high abundances, and found that our approach also effectively predicts metabolic behaviors of host-associated bacteria. In the setting of human health, understanding metabolic changes within bacteria in environments where growth substrate availability is difficult to quantify can have large downstream impacts on our ability to elucidate molecular drivers of disease-associated dysbiosis across the microbiota. Our results indicate that RIPTiDe may have potential to provide understanding of context-specific metabolism of bacteria within complex communities.

**Author Summary:** Transcriptomic analyses of bacteria have become instrumental to our understanding of their responses to changes in their environment. While traditional analyses have been informative, leveraging these datasets within genome-scale metabolic network reconstructions (GENREs) can provide greatly improved context for shifts in pathway utilization and downstream/upstream ramifications for changes in metabolic regulation. Many previous techniques for GENRE transcript integration have focused on creating maximum consensus with input datasets, but these approaches were recently shown to generate less accurate metabolic predictions than a transcript-agnostic method of flux minimization (pFBA), which identifies the most efficient/economic patterns of metabolism given certain growth constraints. Despite this success, growth conditions are not always easily quantifiable and highlights the need for novel platforms that build from these findings. Our new method, RIPTiDe, combines these concepts and utilizes overall minimization of flux weighted by transcriptomic analysis to identify the most energy efficient pathways to achieve growth that include more highly transcribed enzymes, without previous insight into extracellular conditions. Utilizing a well-studied GENRE from *Escherichia coli*, we demonstrate that this new approach correctly predicts patterns of metabolism utilizing a variety of both *in vitro* and *in vivo* transcriptomes. This platform could be important for revealing context-specific bacterial phenotypes in line with governing principles of adaptive evolution, that drive disease manifestation or interactions between microbes.

## Introduction

Metabolic plasticity is critical for bacterial survival in changing environments, and fundamental to stable microbial communities [1, 2]. In the context of human health, it has been shown that certain pathogens adapt their metabolism to their current environment to most effectively colonize a new host [3, 4]. Fully understanding these shifts and their implications may provide opportunities for novel therapeutic strategies. However, describing these situational differences in overall metabolism of an organism have remained difficult to quantify. While genomic analyses of these species reveal catalogues of metabolic capabilities for a species, they do not provide information toward the inter-connected networks of metabolism that invariably occur within the cell. One approach that addresses these limitations is computationally combining genomic data with known biochemical constraints for predicting/modeling cellular physiology with genome-scale metabolic network reconstructions (GENREs). A GENRE is composed of the collection of genes and metabolic reactions associated with the species of interest, representing the totality of known metabolic function that organism is able to employ. This functionality can then be formalized with a mathematical framework and constrained by known biological parameters to allow for simulation of metabolic processes. These powerful discovery platforms have enabled guided genetic engineering efforts, directed hypothesis generation for downstream laboratory testing, and investigation of metabolic responses of bacteria to antibiotic stress [5, 6].

GENREs have also become strong platforms for providing context to multiple forms of high-throughput omics data, allowing for the study of whole-network changes in metabolism due to upstream/downstream shifts in enzyme activity [7]. To this end, advances over the last decade in RNA-Seq technologies have indeed enabled whole bacterial community metatranscriptomic analysis [7]. Conversely, changes identified through other means occasionally can be difficult to attribute to a single organism from complex biological samples (e.g. untargeted metabolomics). However, shotgun transcriptomic analysis followed by strict read mapping protocols allows for the characterization of individual groups of species within these communities of organisms [8, 9]. These large datasets provide informative possibilities for analyses; however holistic (top down) analyses are limited in their ability to make any mechanistic claims outside of correlative results. Against the backdrop of the reductionist (bottom up) approach of GENREs, their combined informative capacity allows for mechanistic insight into the metabolic status and ecological interactions within the microbiome [10].

Numerous approaches to integrating transcriptomic data into GENREs primarily focus on identifying reactions associated with defined cutoffs of gene transcription or change, then maximizing flux through those reactions with many transcripts and minimizing flux through those reactions associated with genes with fewer transcripts [11–21]. While these techniques have produced insights into the influence of transcription on the metabolic state of cells, the largely arbitrary thresholds dividing genes and associated reactions into categories of activity or probability deviates from the continuous nature of mRNA abundance data and can skew results. Several alternative platforms collectively rely on identifying core sets of metabolic functionalities that define the cell-type or condition of interest utilizing omic data, then selecting the most likely pathways or network topology for including these tasks and pruning unused sections of the models [22, 23]. While these approaches have been shown to create models that contain increased amounts of known tissue-specific metabolic pathways, they do not inherently generate fully functional models of metabolism nor are they necessarily reflective of momentary metabolic trends that would be active under specific conditions. Overall, very few platforms have integrated continuous weights to reactions based on transcriptomic data as well as applying these constraints to fully functional metabolism [24–26], but such methods ultimately require high degrees of input data (e.g. fluxomics or differential expression analysis) and rely on placing new bounds on metabolic reactions in relation to the input transcriptome. Of these methods, some have found a measure of success through maximizing correlation between reaction flux and associated transcript abundances [27, 28], yet as previously stated may be a less biologically valid approach as transcript levels are not always predictive of corresponding protein concentrations. Even fewer studies have incorporated this concept that transcription does not necessarily equate with functional enzyme by examining its implications in flux analysis [28], highlighting an opportunity for novel approaches to analyze context-specific metabolic models in relation to continuously assigned reaction weights and stand-alone transcriptomic datasets.

Recently, parsimonious Flux Balance Analysis (pFBA) was developed, which seeks to minimize the flux associated with each reaction in the model while maintaining optimum flux through the objective function [29]. In essence, this approach identifies the least biologically “expensive” usage of an organism’s metabolism to achieve high growth rates. This method is in line with the concept that evolutionary pressures have selected for metabolic states in microbes with minimized cellular cost that maximize growth rate under a variety of environmental conditions [30]. Surprisingly, with a model of *Saccharomyces* metabolism, pFBA alone with constraints on metabolic inputs was found to be able to largely outperform several other methods in predicting metabolic network behavior under various growth conditions despite not utilizing any transcriptomic data for these predictions [31]. Recently, our laboratory has also demonstrated that parsimonious flux analysis, in conjunction with differential expression, can augment predictions of metabolite output with both rat and human GENREs [32]. These studies demonstrate that contextualizing data in a truly biologically meaningful way has proven difficult, highlighting large gaps in our understanding of metabolic regulatory mechanisms. Additionally, recent studies on *Escherichia coli* grown in defined *in vitro* conditions have shown that levels of transcript were only predictive for roughly half of the concentrations of the corresponding enzymes in the cell [33, 34]. In reality, reaction flux is determined by a complex combination of local substrate and enzyme concentrations as well as the kinetics of the reaction itself. Despite this shortcoming, transcript abundances remain an indication of cellular investment into a given metabolic strategy. Furthermore, as the expression of a functional enzyme is not as simple as an on-or-off scenario, we have also implemented an unsupervised procedure to assign weights to reactions based on the distribution inherent to RNA-Seq data [35]. As arbitrary thresholds have been demonstrated to be biased [36], and absolute mRNA abundances were shown to improve GENRE flux predictions when their correlation with reaction activity is maximized [28], we chose to calculate continuous values along the transcript abundance distribution for the coefficient associated with each gene. This approach is unbiased by depth of sequencing across datasets, and it restricts the utility of low-transcription reactions to an optimal pFBA solution yet does not entirely prohibit it. With both of these concepts in mind, we have leveraged the insights gained by pFBA, integrating transcript abundance as a weighting metric to direct the most parsimonious flux solutions toward a state of higher fidelity to *in situ* biology.

We call this combined method RIPTiDe, or Reaction Inclusion by Parsimony and Transcript Distribution. RIPTiDe was then tested on the most well curated GENRE of the highly studied model organism *E. coli* str. K-12 substr. MG1655. Transcriptomic data from multiple distinct *in vitro* growth conditions were compared against each other as well as the base implementation of pFBA. RIPTiDe was able to accurately discern context-specific phenotypes including growth rates that closely match experimentally measured values under the same conditions, as well as gene essentiality predictions in metabolic pathways relevant to the corresponding media. The platform was subsequently tested utilizing *in vivo* metatranscriptomic sequence data from clindamycin-treated mouse cecal content where *E. coli* is the dominant member of the bacterial community [8]. When contrasted against results from the previous analysis, the *in vivo* RIPTiDe-contextualized model simulated significantly slower growth than in aerobic rich media and more central metabolites were produced intracellularly when in competition with other bacterial species. These combined analyses validated our methodology by correctly uncovering context-specific biological trends, and previously unappreciated host-associated behaviors of *E. coli*. Furthermore, this technique could present a powerful tool for the study of the microbiota through the simultaneous unsupervised contextualization of omics data from multiple organisms within the same community.

## Results and Discussion

### Reaction Inclusion by Parsimony and Transcript Distribution (RIPTiDe)

RIPTiDe relies on sequential optimization steps, guided by transcript abundances in order to identify the most efficient usage of metabolism that also reflects an organism’s transcriptional state [29]. The required inputs for RIPTiDe are a GENRE (with an assigned objective) and a dictionary of gene-associated transcript abundances. By first calculating the optimal objective value for the original GENRE, and then constraining any future optimization to approximately this level of flux, RIPTiDe ensures both functional output models as well as the identification of highly efficient metabolic strategies that are consistent with transcriptomic datasets. Subsequently, transcript mapping is reduced to only genes that appear in the genome-scale network reconstruction of interest, therefore not skewing the distribution with data that is uninformative to the current analysis. To generate reaction coefficients (weights) that closely reflect the normalized transcript abundance [37] for their associated genes, we calculated the ratio of transcription for each gene with the maximum abundance in the dataset (Fig. 1A). In line with the overall motivation for RIPTiDe, this calculation results in a weight for all genes that represents each as a fraction of the maximum cellular investment in any one metabolic function, which has been shown to be effective in at least one previous approach for transcriptome integration [25].

**Figure 1.**
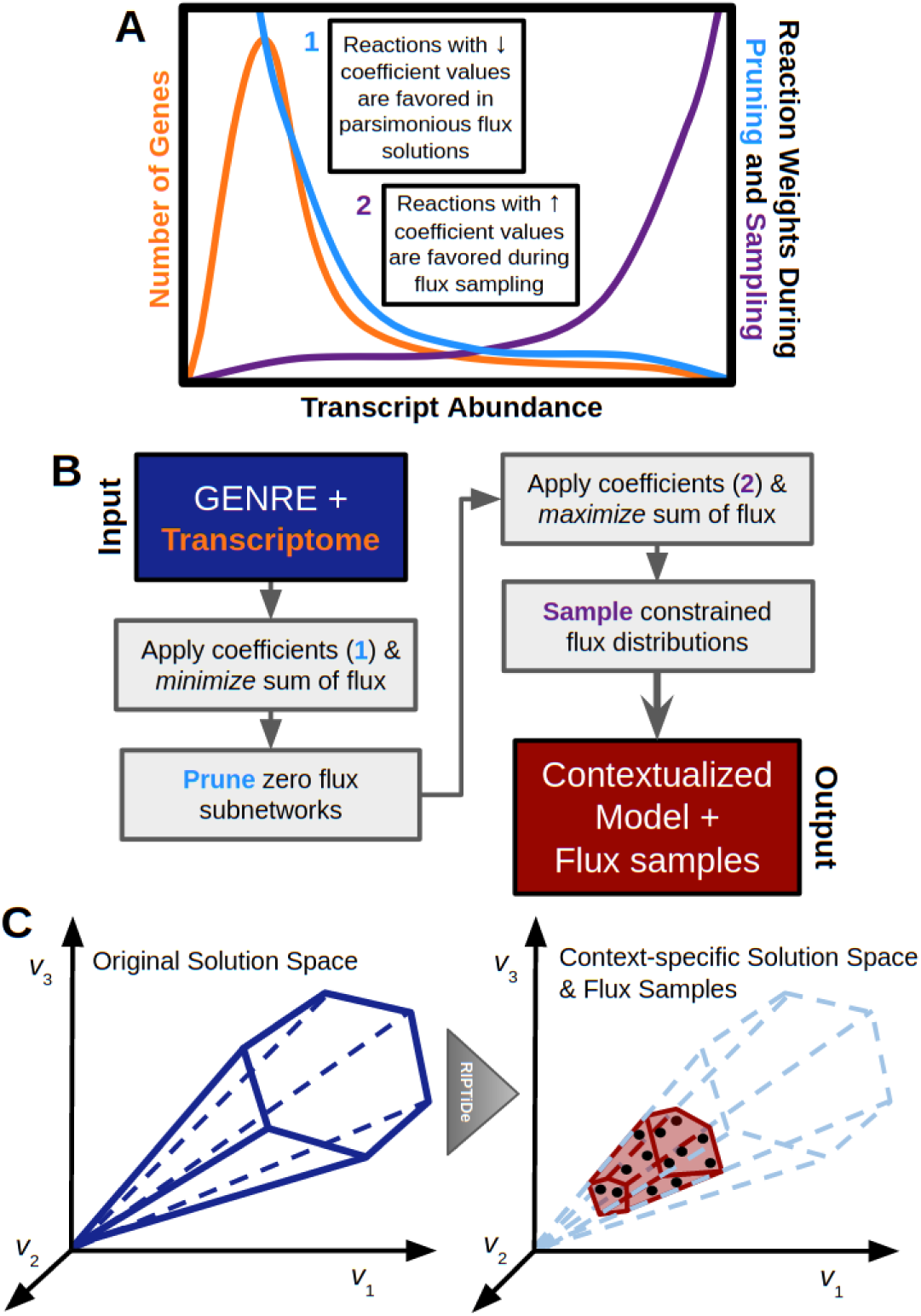
Visual representation of RIPTiDe workflow. **(A)** Example transcript and resultant reaction weight distributions were calculated by RIPTiDe. During the pruning step, reactions for genes that recruit greater abundances of transcript are assigned *smaller* linear coefficients which in turn result in higher likelihood of usage in an overall flux minimization objective. Alternatively, during the subsequent the flux sampling step, the remaining reactions associated with higher transcription are assigned *larger* linear coefficients leading to increased possible flux ranges. **(B)** Schematic outlining computation performed during each step of RIPTiDe. The platform requires two input data structures: a genome-scale network reconstruction and transcript abundances associated with genes. During initialization transcript abundances for genes are transferred to their corresponding reaction, allowing coefficients to be assigned. Each reversible reaction is now also separated into pairs of irreversible reactions. Applying principles of parsimonious FBA to the constrained model, a minimum sum of fluxes optimization is performed with respect to a predefined minimum flux through the cellular objective. Reactions that no longer carry flux in this state are pruned, and flux sampling is performed on the intermediate model to determine context-specific bounds for the remaining reactions. **(C)** Pictorial representation of the impact of RIPTiDe on the original reconstruction, converting it to a context-specific model of metabolism with a parsimonious metabolic solution space with respect to the given transcriptomic data.

The process results in a distribution of linear coefficients that is >0.0 and ≦ 1.0; a higher transcript abundance corresponds with a lower linear coefficient and therefore a larger probability of being utilized in the final metabolic model, given that RIPTiDe first minimizes the sum of linear coefficients and fluxes. Additionally, in the cases where multiple genes contribute to the same reaction (i.e. isozymes or multimers), then the lowest coefficient (highest transcript abundance) among the group is chosen as it reflects in what processes the cell has invested energy. Another approach is have integrated overall GENRE flux state limited by whole-cell transcriptomic analyses to more directly predict metabolic capacities [25]; however, this approach did not leverage additional constraints on total flux, as done here through a pFBA assumption, to maximize cellular economy as a reflection of omics datasets. By calculating a continuous scale of linear coefficients based on absolute transcript abundances, RIPTiDe removes the need for different cutoffs between datasets of variable sequencing depth and reduces the requirement for manual intervention in the generation of functional context-specific models.

As with pFBA, reversible reactions are split into pairs of irreversible reactions carrying non-negative flux. Coefficients are subsequently transferred to the corresponding metabolic reactions for each gene. Reactions are then assembled into a single new equation which is set as the cellular objective where the magnitude of the linear coefficient drives the amount of flux carried by each reaction. Reactions without genes are assigned the median score to not be incorrectly penalized, nor overly biased to their inclusion. Larger coefficients correspond to the reaction receiving a larger weight per unit of flux during optimal growth, which would therefore be favored less when minimizing the overall sum of weights times flux for a given the objective function. In this way, we were able to minimize overall flux subject to the transcript-guided linear coefficients added by RIPTiDe. Within the transcript abundance distribution, progressively smaller linear coefficients are assigned to genes with increased transcription (Fig. 1A). With an additional linear optimization, reactions with no flux are pruned from the network. As no arbitrary cutoffs are introduced during these steps, this approach is equally tractable between datasets of variable sequencing depths.

The scope of feasible solution states in a GENRE typically out-scales the number of realistic metabolic states of the cell by a large magnitude [38]. Placing further constraints on a model that reflect likely metabolism allows for improved predictions, but only partially mitigates the potential for biologically meaningless solutions to be reported. Sampling of feasible flux distributions reveals how the applied constraints limit the possible metabolic states and ensure we do not rely on any single state that may be physiologically less likely [39, 40]. Each sampled flux distribution represents a hypothesis for how metabolism is truly functioning given the constraints placed on the system. Integrating a flux sampling technique into RIPTiDe, we were able to further leverage our transcript-based constraints to maximize the context-specificity of resultant models. Following assignment of linear coefficients to metabolic reactions, we proceeded through the linear optimization steps at the core of RIPTiDe as illustrated in Fig. 1B. Of note, after reaction pruning we assign weights to the reaction fluxes directly proportional to transcript abundance (inverse of how the weights are applied before pruning) because we want to promote flux through the remaining reactions that are associated with high transcript values during the flux sampling analysis. In order to be quickly and uniformly sample this highly-constrained solution space, we utilized the sampler gapsplit which is tuned to maximize efficiency per sample under this type of condition [41]. Transcript abundances were used to guide the optimization and flux sampling in order to identify the most economic, likely usage of metabolism with the given cellular investment in transcription. A more detailed description of the steps in RIPTiDe are as follows:

1. Near optimal flux through the objective function is set as a constraint (≧80%, Table S1).
2. Linear coefficients are assigned as constraints to each reaction based on transcriptomic abundance of the associated gene-enzyme assignments. In cases where multiple gene products contribute to the same reaction, the lowest coefficient/largest weight among them is selected.
3. Total sum of weights times fluxes in the model is minimized subject to previously outlined constraints.
4. Standard flux balance analysis then identifies inactivated subnetworks (zero flux) which are subsequently removed from the model. Orphaned genes and metabolites are then pruned.
5. Inverse linear coefficients within the original provided range are calculated and assigned for each reaction. Flux sampling is performed on the constrained and reduced model to find most likely pathway utilization patterns while maintaining 80% of the objective function.

The resultant pruned model has a dramatically more focused, biologically feasible, metabolic solution space that has been adequately sampled for feasible metabolic states (Fig. 1C). Unlike many previous approaches, models produced by RIPTiDe are able to achieve flux through their given objectives by default without any reintroduction of deactivated reactions. This characteristic is critical as the principle of optimum cellular economy with respect to the transcript data is never broken throughout the contextualization process. In addition, maintaining flux through a given objective function allows for predictions on potential targets for therapeutic interventions, such as essential genes. Spearman correlation is then performed between median absolute flux values for all reactions found during sampling and their associated levels of transcription to identify the level of concordance of context-specific metabolic states with the measured transcriptome (Fig. S3A). Differences between median absolute flux values and transcript abundances highlight reactions that are potentially subject to high post-transcriptional regulation [42]. Furthermore, we have also integrated functionality to instead utilize one or more metabolic tasks (single reactions) as constraints which must achieve positive flux in order to accommodate larger-scale models where a single defined cellular objective may be a biologically invalid assumption (e.g. human tissue). For a more detailed description of the algorithm, refer to the Methods section.

### Application of RIPTiDe to an Example Model of Bacterial Metabolism

To test RIPTiDe, we created a small, flux-consistent metabolic network reconstruction with two compartments and generation of ATP (in arbitrary units) as the cellular objective function. The model has two separate means of accomplishing this goal; either through glycolysis where glucose is the primary substrate, or Stickland fermentation which requires pairs of amino acids in coupled redox reactions to create energy [43]. In nature, fewer ATP are generated per unit of substrate in Stickland fermentation compared to glycolysis [44]. This phenomenon was replicated in the example model as fewer reactions are required to generate ATP from glucose, making this pathway more energetically favorable (Table S2). Utilizing the base implementation of pFBA available within COBRApy to quantify which pathway was the more parsimonious solution, we ensured that glycolysis was actually the most efficient (lower sum of flux with fewer enzymes used) for optimal ATP generation (Fig. 2A). Critically, this ATP generation was performed under the conditions where all possible extracellular substrates were provided in excess to the model as to not bias results only on the basis of resource availability.

**Figure 2.**
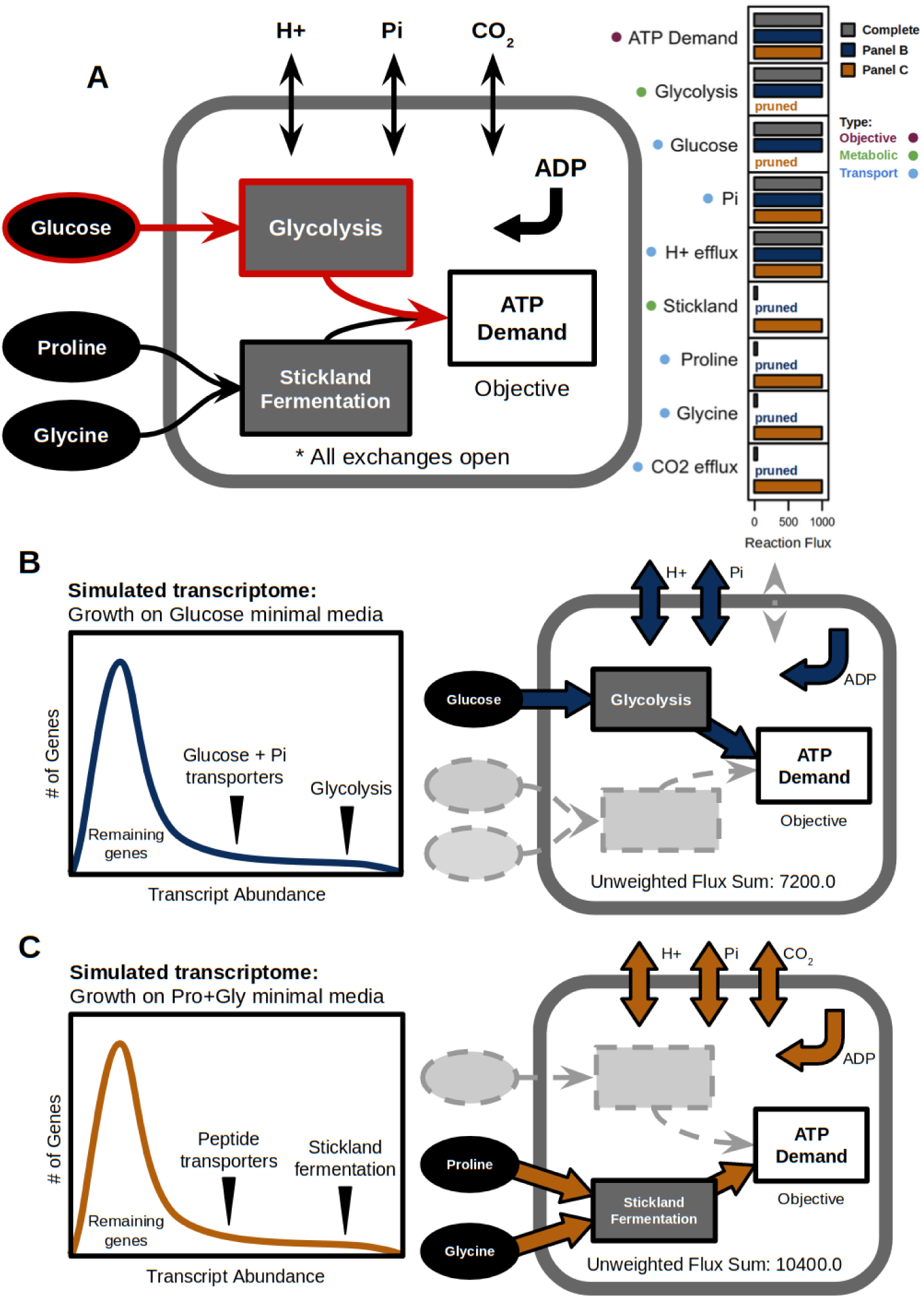
Example model of bacterial metabolism supports the utility of RIPTiDe for identifying most likely context-specific strategies. **(A)** General topology of simplified GENRE that consists of 16 reactions and 14 metabolites in total (Table S2). The objective of ATP generation can be achieved through two separate means: either the catabolism of glucose via glycolysis or through the paired fermentation of proline and glycine. Glycolysis is the more parsimonious pathway for generating ATP, which is reflected in the pFBA solution fluxes to the right of the diagram (outlined in red). **(B)** RIPTiDe is able to correctly identify a likely route of flux when provided a transcriptome. **(C)** When provided with transcriptomic evidence, RIPTiDe is able to identify a less parsimonious, but more concordant solution (Lower *weighted* sum of fluxes). Despite very low transcription, RIPTiDe still identifies the necessity for carbon dioxide efflux during fermentation to maintain mass balance.

As a means to determine the context specificity of RIPTiDe output models, we then generated two distinct simulated transcript abundance distributions that each reflected elevated transcription of a single pathway (Table S2). In Fig. 2B, RIPTiDe was provided with a transcriptome in which genes for glycolysis and related reactions were more highly transcribed. The contextualized model in this case also utilized glycolysis as the only means of generating ATP, directly matching those results obtained from pFBA. Alternatively, as illustrated in Fig. 2C, when genes associated with Stickland fermentation and peptide substrate transporters were more highly transcribed, RIPTiDe was able to discern that the less directly parsimonious pathway should be utilized instead when provided sufficient transcriptomic evidence. In each case, reactions related to the opposite pathway were correctly pruned while generating the final context-specific model. Furthermore, flux samples for the cellular objective from each contextualized example model revealed that significantly more ATP could be produced when employing glycolysis (Table S2), further supporting the energetic favorability of this pathway. These combined results confirmed that RIPTiDe was indeed able to use a less parsimonious pathway when necessary to better reflect the associated transcriptomic dataset. We also made the same topological assessment following Stickland fermentation-associated transcriptome integration using two other relatively recent approaches, CORDA and RegrEx [13, 27]. Neither algorithm yielded the same result as RIPTiDe, and both incorrectly retained glucose and its associated transporter in their resultant models (Table S2). These results further indicate that RIPTiDe is well suited to identify metabolic capabilities which are most likely to be utilized given cellular economy and transcriptomic evidence.

### RIPTiDe Performance with a Curated Model of *E. coli* Metabolism

To perform a thorough evaluation of our approach, we utilized the genome-scale metabolic network reconstruction iJO1366 (2583 reactions, 1805 metabolites, 3 compartments), for the model organism *E. coli* strain K-12 substr. MG1655 [45]. This GENRE has been the subject of extensive manual curation and experimental validation, making it the best possible candidate for examining new approaches in constraint-based modeling [46]. Leveraging these years of research, we utilized this well described system to determine the biological validity of predicted context-specific differences returned by RIPTiDe. To accomplish this goal, we performed a comparative analysis using three separate transcriptomic sequencing datasets, each derived from well characterized *in vitro* media conditions including aerobic LB rich media, aerobic M9+glucose defined minimal media, and anaerobic M9+glucose defined minimal media. Each dataset was collected at approximately the same mid-exponential phase of growth [47, 48], and contextualization of iJO1366 with RIPTiDe took only seconds to complete (Fig. S3B). These data were chosen due to the large amount of characterization each has received, allowing our predictions to be more readily supported compared to assessing metabolic differences possibly seen across numerous less well explored conditions. Additionally, *E. coli* is a facultative anaerobe with known differences in metabolism between these lifestyles that we hoped to capture with RIPTiDe [49].

We first set a performance baseline using implementations of the distinct methodologies for contextualizing omics data with GENREs mentioned previously; strict integration to corresponding reaction flux levels versus complete context-specific model generation. For equal comparison between algorithms, these analyses were all performed utilizing the transcriptome collected from *E. coli* K-12 growing in aerobic LB rich media. We elected to assess an array of distinct GENRE+transcriptome integration algorithms; including GIMME, iMAT, CORDA, MBA, and RegrEx [11–13,20,27]. Of note, GIMME and CORDA require the resulting model to meet an objective function while iMAT, MBA, and RegrEx maximizes the number of reactions present in the model based on a defined threshold, and therefore do not necessarily meet a cellular objective. Additionally GIMME, iMAT, CORDA and MBA all require abundance cutoffs for selectively including reactions associated with genes that meet predetermined levels of transcription, while RegrEx does not. Instead, RegrEx attempts to maximize correlation between model activity and transcript levels through least squares regression, which does not completely reflect functional enzyme concentration nor focus on obtaining a functional model [33, 34]. In several of the algorithms, reactions necessary for an objective function do not necessarily have associated GPR rules, meaning they will be considered inactive and pruned from the network. Additionally, it is important to note that each of these algorithms obtain slightly varied results and may be better suited for their original analysis objectives. For all platforms, we utilized the recommended threshold values published alongside each method as well as allowing for all available substrates to be taken in by the system. With these settings only GIMME, CORDA and RIPTiDe returned models that were able to achieve flux through the biomass objective reaction (Table S3). GIMME and CORDA yielded models containing 2055 and 1702 reactions respectively, both without a substantial shift in growth rate. This level of pruning did not substantially narrow the possible compensatory pathways for achieving growth which can be studied easily, and also does not necessarily reflect a single context-specific metabolic strategy. Alternatively, RIPTiDe created a model which only contained 567 reactions and a large decrease in flux through the biomass reaction, consistent with the shift in doubling times between rich and minimal media. This point was even better represented in a comparison of context-specific gene essentiality which revealed large differences across platforms that yielded functional models (Table S3). From genes that are included in iJO1366, we limited our analysis to only focus on those also found to be experimentally essential in a single gene deletion screen of *E. coli* K-12 growing in LB rich media [50]. Of the genes from this list that were present in the RIPTiDe-generated model, all were found to be essential for biomass generation. Conversely, only 72% and 65% of genes from GIMME and CORDA-generated models that were experimentally essential were also essential *in silico*. Only after altering extracellular metabolite availability to reflect LB media components are all genes found to be essential across all platforms (Table S3). These results support the necessity for techniques that do not require such a high degree of information at the outset.

### Growth Substrate Uptake Predictions Using RIPTiDe Reflect Experimental Measurements

To evaluate how closely RIPTiDe-based predictions reflect predictions when media conditions are known, we performed NMDS ordination analysis of Bray-Curtis dissimilarities between common exchange reaction flux distribution samples from iJO1366 with and without constraints on available substrates compared with RIPTiDe-generated flux samples with no defined media (Fig. 3A). For these comparisons, RIPTiDe was applied to iJO1366 with no additional constraints applied utilizing the transcriptome from growth in M9+glucose defined minimal media. As expected, flux samples from the model with only constraints for near maximal growth are significantly different (*p*-value = 0.001) from each other group as it has access to all possible growth substrates which combined allow for faster growth and distinct patterns of metabolism. A striking relationship emerged when RIPTiDe flux distributions were compared with those from media-constrained models, revealing that these groups were highly correlated. Importantly, no difference (*p*-value > 0.05) was found when RIPTiDe was applied to the model with M9+glucose media constraints contrasted against results from RIPTiDe-generated flux samples without media constraints (Table S4). A similar trend was also observed with pFBA integrating two additional transcriptomic datasets from *E. coli* across distinct media conditions (Fig. S5). These data indicated that RIPTiDe-based predictions for an organism’s interaction with its environment (exchange fluxes) were analogous to those made when extracellular metabolic conditions were known. With these results in mind, for all subsequent work we took an unsupervised approach by allowing for unpenalized flow of all extracellular metabolites in and out of the system (exchange reaction bounds set to ±1000), otherwise referred to as “complete” media. This approach allowed for inferences about the most likely growth substrates for a contextualized GENRE since only those that carry flux following RIPTiDe remain, which is imperative to accurately describing context-specific metabolic patterns in complex extracellular environments. We performed this inference in aerobic LB rich media as well as aerobic M9+glucose defined minimal media (Table S7). Growth substrates were defined as those extracellular metabolites which had a negative median flux through the associated exchange reaction following flux sampling of the context-specific models. This analysis in LB rich media revealed that 54 metabolites (76.1% of the total remaining exchange reactions after pruning) were generally imported by the RIPTiDe-generated model and used for growth. Among these substrates, most have been documented as constituents of LB including numerous amino acids which were predicted as the primary *in silico* carbon sources. This finding reflects known patterns of *E. coli* physiology in LB as the media is void of sugars that are fermentable by the bacterium, and primarily utilizes several amino acids for growth [51, 54]. Additionally, glucosamine and glycerol 3-phosphate were predicted carbon sources in rich media, both of which have been shown to be commonly recycled components of the *E. coli* cell wall during growth in LB [60, 78]. When contrasted against those growth substrates RIPTiDe predicted with the M9+glucose transcriptome, glucose and fructose were instead included as carbon sources which appropriately tracked with known *E. coli* metabolic behaviors [51–54]. Conversely both models produced by GIMME and CORDA using the LB-associated transcriptome were unable to perform similarly; with GIMME predicting net uptake of mainly trimethylamine, and CORDA predicting both butyrate and indole as primary sources of carbon (Table S7). None of these metabolites are found in LB and *E. coli* lacks the cellular machinery to produce them for eventual recycling, making their availability *in vitro* improbable. As an additional finding, the median flux values predicted by RIPTiDe were within realistic ranges for rates of metabolite import and not nearing the arbitrary high bounds (±1000) as was the case with GIMME and CORDA-produced models. These results collectively indicated that RIPTiDe is also able to approximate the media condition from which the transcriptome was collected. This feature will likely be especially useful when growth conditions are much more difficult to quantify, as in environmental samples or *in vivo*.

**Figure 3.**
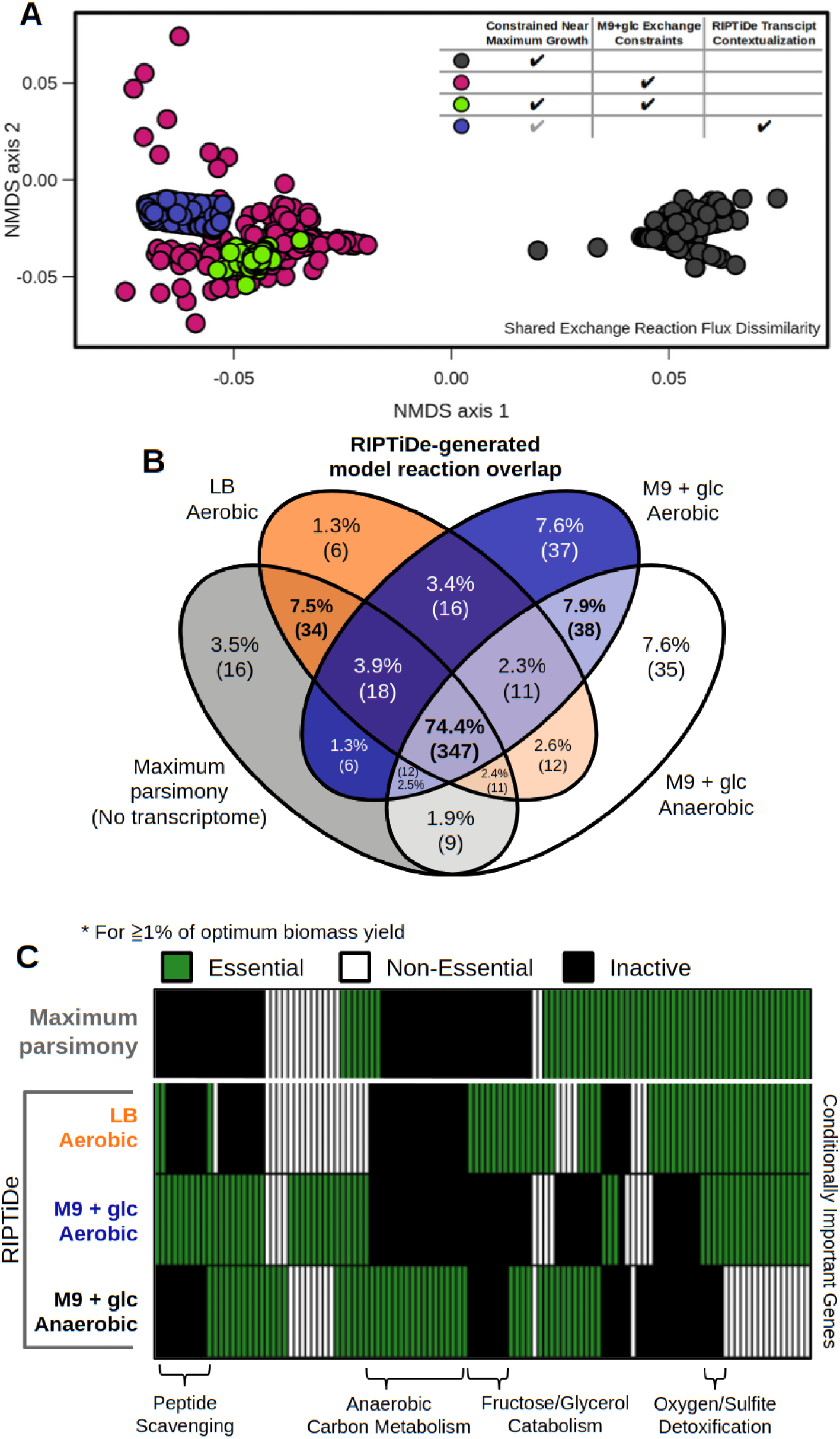
RIPTiDe identifies established biological differences in *E. coli* str. K-12 substr. MG1655 across *in vitro* growth conditions. Without increased user supervision, RIPTiDe correctly predicts behavior and context-specific pathways for *E. coli* str. K-12 substr. MG1655 (iJO1366), while simultaneously producing functional models that can be characterized phenotypically. **(A)** NMDS ordination of Bray-Curtis dissimilarities between exchange reaction flux samples for each version of iJO1366. Table legend indicates flux constraints placed on iJO1366 prior to flux sampling. Flux samples from RIPTiDe transcriptome contextualization without exchange constraints are not significantly different from those where media condition was set *a priori* (*p*-value > 0.05). The gray check denotes that the max growth constraint is inherently integrated into the RIPTiDe workflow. Significant differences evaluated by PERMANOVA. **(B)** Comparison of metabolic reactions included among RIPTiDe-contextualized transcriptomes with iJO1366. The majority of reactions (71.5%) are conserved across models within a central set of pathways. All exchange reaction bounds set ±1000 prior to contextualization. **(C)** Comparison of importance (essentiality) for conserved genes across pruned RIPTiDe models. 105 core essential genes were identified across all groups (Table S3). Hierarchical clustering reveals context-specific pathway essentially, labeled across the bottom axis, based on the environment in which the bacterium is growing. All exchanges set to ±1000 prior to contextualization.

### Context-specific Differences in *Escherichia coli* Biology Uncovered by RIPTiDe

We next performed RIPTiDe-based analyses of iJO1366, utilizing all three transcriptomic datasets described previously. As a standard of comparison, we also created a simulated transcriptome where all transcript abundances were identical, forcing RIPTiDe to assign maximum importance to all reactions and establish the most “economic” overall flux distribution that achieves maximum growth (referred to as Maximum Parsimony), and also constrained to achieve at least 80% of the optimal objective function value. We first assessed the impact that each transcriptome had on the components of each model created by RIPTiDe. Different topologies of reaction inclusion were found among each of the contextualized models; however none were completely unique and a core set of metabolic reactions were necessary for optimal growth across all conditions (347 reactions; Fig. 3B). The next largest fraction of overlap was between those models generated using transcriptomes from *E. coli* grown in M9 minimal media (aerobic & anaerobic), which supported that RIPTiDe was able to identify general biological trends that are conserved between similar media conditions. Interestingly, the next largest group was the intersection between Maximum Parsimony and LB rich media, which may reflect that growth in an environment with surplus growth nutrients is partially analogous to the artificial state of minimal cellular investment. These distinctions in network topology were further crystallized by conditional gene essentiality between the RIPTiDe-contextualized models (necessary for at least 1% of optimum growth). It is important to note that genes found to be essential during these analyses do not necessarily correspond with survival of *E. coli*, only that these genes may be critical to the optimal metabolic strategy under the associated conditions [55]. Future work may seek to elucidate the individual contribution of these genes to overall metabolic strategies across respective media conditions. Although 105 genes were essential all four contextualized models, in addition to the 152 essential genes in the uncontextualized GENRE that were also essential in all contextualized models (Table S5), those genes that were context-specific often related to pathways of biological significance to their respective media conditions (Fig. 3C). As a corollary to creating a smaller metabolic network than those using transcriptomes contextualized by RIPTiDe (by an average of 9%), fewer genes were distinctly essential in the model resulting from Maximum Parsimony (5 genes) compared to any of the RIPTiDe-contextualized *in vitro* transcriptomes (8, 15, and 20 genes respectively; Table S5). Interestingly, the contextualized models on average shared larger ratios of components (approximately 7% reactions) with each other than with Maximum Parsimony, supporting that RIPTiDe selects more distinct patterns of metabolism compared to Maximum Parsimony alone (Fig. 3B). We also calculated the relative doubling time for each of the contextualized models from the objective values associated with the biomass objective function (Fig. S1) [56]. As expected, the base model with all open exchanges and no additional constraints displayed the fastest doubling time. However, the growth rates for models associated with *in vitro* transcriptomic data each decrease in the expected order reflecting experimentally measured results [51, 57]; LB (aerobic) → M9+glucose (aerobic) → M9+glucose (anaerobic). These results collectively suggested that the RIPTiDe-contextualized GENREs not only topologically reflected known biological trends, but also phenotypically mimicked experimentally measured behaviors.

We subsequently focused our analysis on assessing known differences in core metabolism across transcriptomes in RIPTiDe-contextualized models. Growth conditions are known to affect which forms of metabolism are possible or favored at a given time, driving the cell to utilize alternative pathways for replenishing it’s cellular currency. One such anabolic circuit in Enterobacteriaceae is the small transhydrogenase pathway which consists of two distinct enzymes, contributing to NADPH homeostasis within the cell [58]. It has been previously demonstrated in *E. coli* str. K-12 that transhydrogenase functionality is critical for normal growth in M9 minimal media, but dispensable in LB rich media [59]. Strikingly, the RIPTiDe-contextualized model in aerobic M9 minimal media indeed obtained the majority of cellular NADPH from this mechanism (Fig. 4A). Alternatively, both gene-associated reactions were pruned when the LB media-associated transcriptome was integrated with RIPTiDe. Instead in rich media, the contextualized model generated the bulk of its NADPH via glycerol-3-phosphate dehydrogenase (Fig. 4B). This enzyme is a major contributor to the bacterial electron transport chain and known to be active in *E. coli* during fermentation and growth in aerobic LB broth [54, 60]. Importantly we found that pntA and pntB, the genes for NADP transhydrogenase subunits, were significantly upregulated in M9 minimal media and downregulated in rich media (Table S5).

**Figure 4.**
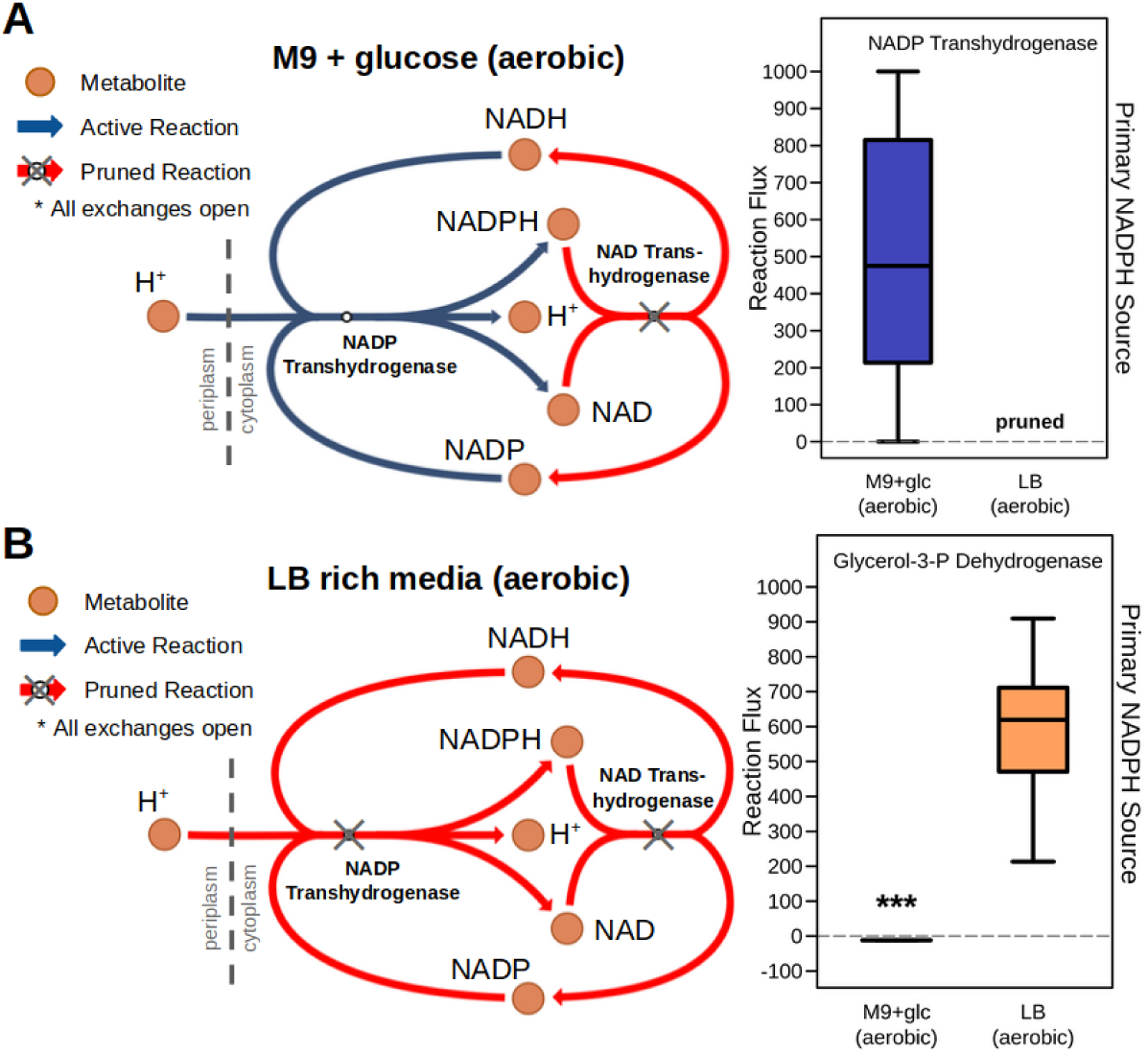
Cellular sources of NADPH in RIPTiDe contextualized models reflect known biological differences in *E. coli* across media conditions. Shown in each panel is reaction pruning of the small transhydrogenase circuit found in *E. coli*, and flux sampling results of NADPH sources from RIPTiDe using the iJO1366 GENRE with contextualized transcriptomic abundances from *E. coli* K-12 MG1655 grown in **(A)** M9+glucose minimal media, and **(B)** LB rich media. This mechanism for restoring NADPH balance is known to be essential for *E. coli* growth in M9 minimal media but dispensable in LB, which is correctly selected by the unsupervised network pruning from RIPTiDe. Significant difference in flux levels determined with Wilcoxon signed-rank test (*p*-value << 0.001); pathway maps generated using Escher [77].

Alternatively, glycerol-3-phosphate dehydrogenase is transcribed relatively evenly in both conditions but contributes to a parsimonious flux solution. This result reinforced that RIPTiDe has the ability to correctly deviate from the most parsimonious flux states when sufficient evidence is presented. These combined findings signified that RIPTiDe is able to capture previously measured, context-specific behaviors with only transcriptomic characterization and no manipulation of *in silico* media conditions.

### Contextualization of *in vivo* Metatranscriptomic Data Reflects Host-Associated Metabolism

Finally, we sought to test the utility of RIPTiDe for capturing metabolic behavior of bacteria from within host-associated communities of microorganisms. Computational analysis of a discrete bacterial species within a complex environment as a member of a larger microbial community has been historically difficult [61]. To address this problem, we utilized *in vivo* metatranscriptomic abundance data collected from the cecum of wildtype C57Bl6 mice. These data were chosen because the specific antibiotic pretreatment (intraperitoneal injection of clindamycin) resulted in a bacterial community composed of >90% *E. coli* [8]. To begin the analysis, we first made certain that the *in vivo* transcript abundance distribution reflected the same negative binomial type as those derived from *in vitro* transcriptomes mapped to the same *E. coli* K-12 MG1655 genome (Fig. 5A). This result demonstrated that with sufficient sequencing depth, a species with a GENRE may be analyzed with RIPTiDe using data collected from among a diverse collection of microbes. After contextualization of this data, the context-specific flux values during sampling were correlated with assigned linear coefficients based on transcript abundances (Fig. 5A & Fig. S2). This result supported that the transcriptomic data was indeed informative, but not overly deterministic, of optimal metabolic states through RIPTiDe. We then assessed how distinctive the flux distributions were of those metabolic reactions which were shared between context-specific models. Cross-referencing each contextualized model to identify shared reactions, we then performed principle component analysis using Bray-Curtis dissimilarity for those flux samples corresponding with this core metabolic subnetwork (Fig. 5B). This analysis revealed not only significant differences between all of the metabolic profiles identified using the transcriptomic data from *E. coli* across the four conditions (*p*-value < 0.001), but more specifically between those bacteria grown in M9 minimal media and those found using LB rich media or *in vivo* transcriptomes (*p*-value << 0.001).

**Figure 5.**
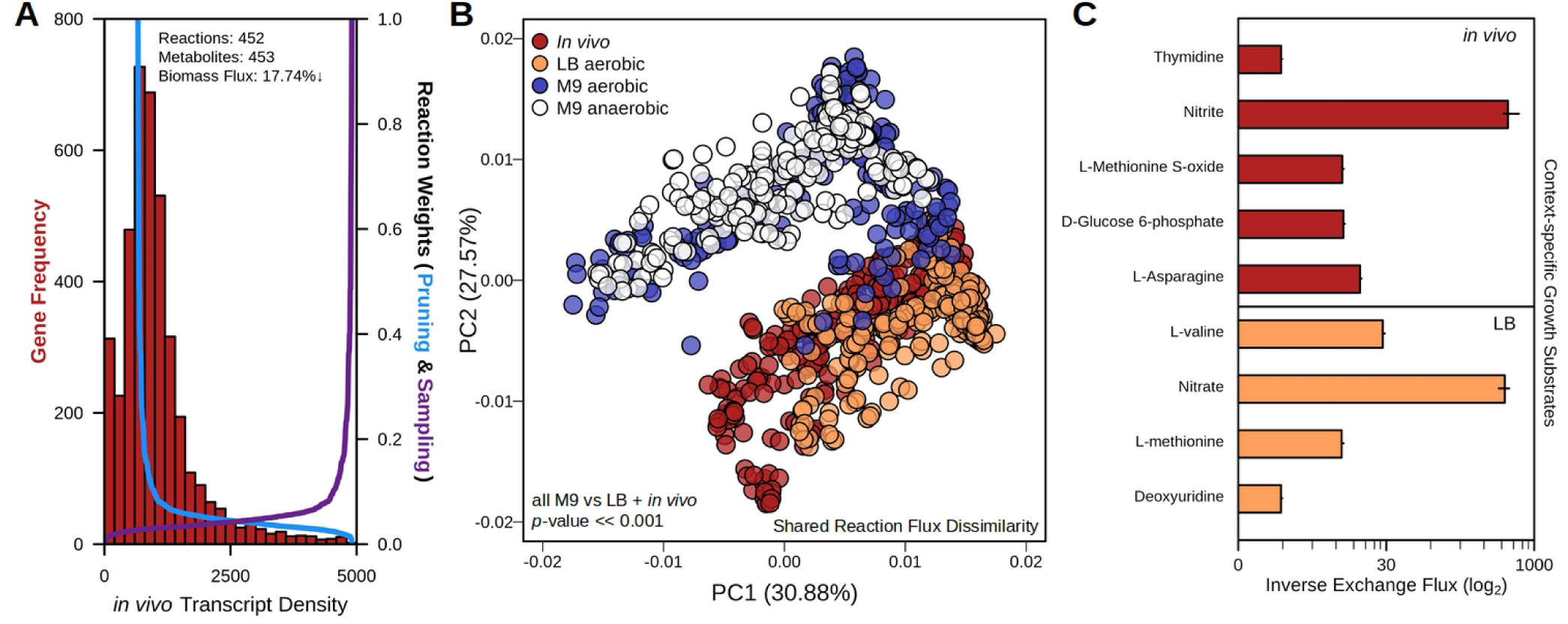
RIPTiDe reveals differential host-associated, metabolism utilizing *in vivo* metatranscriptomic data. Transcriptomic reads attributable to *E. coli* were extracted from a metatranscriptomic dataset sequenced from the cecum of mice in which *E. coli* is the most highly abundant community member [8]. **(A)** *In vivo* transcript abundance of reads recruited to the *E. coli* K-12 MG-1655 genome (left y-axis) and the weights assigned by RIPTiDe to the associated metabolic reactions during each step (right y-axis). Metrics listed at the top of the plotting area reflect the resultant context-specific model compared to the complete iJO1366 model. **(B)** PCoA ordination of Bray-Curtis dissimilarities between flux distributions among reactions of each contextualized iJO1366 model either from *in vivo* or LB rich media (aerobic, *in vitro*) conditions. Significance was calculated by PERMANOVA. **(C)** Metabolite substrates of exchange reactions exclusive to the contextualized models shown. Inverse, normalized flux was calculated by dividing each flux sample by the associated flux through biomass in the same distribution, then multiplied by the overall median flux through biomass across all conditions analyzed. Median and interquartile ranges are displayed.

These results implied distinct metabolic strategies necessary for growth during resource scarcity or surplus, and these patterns were captured by RIPTiDe. Although similar, no group of flux distributions entirely overlapped another and indicated that although numerous reactions and metabolites are shared between all context-specific models, they are utilized significantly differently to manage growth across media conditions and supported differential metabolism is likely necessary to cope with distinct environments.

Next, we contrasted the RIPTiDe models generated using *in vivo* or LB rich media transcriptomes as they were more similar during unsupervised learning analysis. Aside from one environment being highly aerobic and the other largely anaerobic, previous research on *E. coli* physiology during growth in liquid LB concluded that the bacterium utilizes amino acids that are simple to metabolize for growth early on, then switches to other substrates in a diauxic-like growth pattern. LB broth has also been shown to have only modest levels of usable carbohydrate growth substrates, increasing the importance of amino acid catabolism for *E. coli* [51]. Additionally, *in vivo* amino acids and simple sugars are contested resources in the cecum as the majority are already absorbed by the host or used by other microbes upstream in the GI tract, making these growth conditions extremely different [62]. As an analogy to differential growth substrates, we looked to assess the presence of context-specific exchange reactions. We identified 4 exchange reactions that were present in the LB-associated model but not *in vivo*, and 5 of these were peptides or peptide conjugates (Fig. 5C). Carbohydrate growth substrates have been demonstrated to be at very low concentrations in LB media. Additionally, *E. coli* has also been shown to preferentially utilize amino acids to generate energy during early exponential phase [51]. Indeed, upon assessing the measured changes in the *in vivo* concentrations of the highlighted metabolites they were reduced in the context where *E. coli* dominated the environment, possibly due to consumption by the microbe (Fig. S2). Additionally, the exchange for glucose-6-phosphate was also only found in *in vivo*-contextualized models, reinforcing that carbohydrates may be less available for growth by *E. coli* compared to amino acid substrates in LB media [51]. Continuing this analysis now with shared intracellular reactions we utilized AUC-Random Forest [63] on the sampled flux distributions for each reaction to determine those metabolic functionalities that most distinguish the two models (Fig. S4). Among other findings, machine learning revealed consistent differences in flux ranges for both flavodoxin and thioredoxin reductases in aerobic LB culture, but not *in vivo*. These enzymes are important for dealing with oxidative stress which would be much more prevalent in the broth culture than in the mostly anoxic environment of the mammalian gut [64]. Putrescine transport also distinguishes *in vitro* and *in vivo* flux levels, which can be converted to succinate and may be an adaptation for survival in the gut where polyamines (like putrescine) are at elevated concentrations [65]. Altogether, our results support that RIPTiDe is also a strong platform for the analysis of bacterial GENRE behavior in the context of complex microbial communities through the use of metatranscriptomic data.

## Conclusions

In order to survive in a given environment, an organism must express the appropriate forms of metabolism to cope with changing conditions. Production of transcript and protein is metabolically expensive, and over-production likely puts the cell at risk if further perturbation is placed on the system [66]. Although recent findings suggest that levels of transcript for a given enzyme product do not always directly correlate with the amount of functional protein present in the cell at a given time [33], these values may be considered a signature of cellular “investment” and therefore may still be useful in interrogation of metabolism. Furthermore, understanding the quantity of even a small number of functional enzymes remains extremely challenging, which negatively impacts our ability to grasp which and why resources are used by the cell [67]. We should then consider transcriptomic abundance data as signatures of cellular energy expenditure where high transcript levels for a particular gene are an indication of large cellular investment related to that particular function, but are not particularly deterministic of the associated phenotype. Parsimonious Flux Balance Analysis approaches the idea of maximizing cellular economy, and we have built on this idea by employing RNA-Seq data to inform where the cell is most likely spending its energy.

We have demonstrated empirically that our algorithm correctly calculated both *in vitro* and *in vivo* metabolic phenotypes with reasonably high accuracy. In addition to these predictions, our method does not require prior knowledge of active metabolic subnetworks which can preclude work with understudied or uncultured species that cannot or have not been grown or characterized in isolation. Furthermore, the ability to accurately capture the context-specific metabolism of bacteria when media conditions are not easily defined will be an important component for the discovery of metabolic behaviors of microbes in complex environments or novel host-microbe metabolic interactions. Some possible limitations of RIPTiDe do exist however. First, as RIPTiDe relies on the distribution associated with shotgun transcriptomic sequencing, it is less suited for other types of omic data in its current form [68]. RNA-Seq reads are easily attributable to single species within the microbiota due to strict genomic mapping techniques and are measured within a single sequencing assay, however with sufficient consideration to reaction weights given other data types RIPTiDe’s functionality could be expanded. Second, by condensing to such a small metabolic network “solution space”, the models generated by RIPTiDe are meant to represent the most probable context-specific “snapshot” of likely metabolism. To counteract potential overfitting to a single metabolic state for more generalizable downstream models, we created options for manually setting a maximum weight that coefficients are allowed as well as providing lists of core functionality that must remain in the context-specific model created by RIPTiDe. In spite of these concerns, we have shown this algorithm to be a strong approach for contextualization of genome-scale metabolic network reconstructions for both *in vitro* and *in vivo* conditions, producing models that closely mirror known biological phenotypes. Additionally, RIPTiDe run time scales linearly with GENRE size and compartmentalization (Fig. S3B), demonstrating that this technique may also prove effective for generating tissue-specific models with GENREs from larger eukaryotes. The implementation of RIPTiDe rewards divergence from the most numerically parsimonious state to one that is more reflective of cellular investment, and in turn may be more illustrative of *in situ* biology. This approach also allows for inferences to be made about substrate preferences or utilization, even in the context of unknown or highly enriched media conditions. In summary, RIPTiDe approaches transcriptomic data integration uniquely from previous techniques and provides a potentially more biologically relevant platform for considering the cellular behavior in complex extracellular metabolic environments.

## Methods

### Parsimonious Flux Balance Analysis Adaptation

RIPTiDe was implemented within COBRApy [69] and is freely available with installation instructions at https://github.com/mjenior/riptide. Optimal flux through the objective function was set as a constraint with a 20% range of allowed variation (based on 80% fraction of optimal objective function value). Following sensitivity testing, this level was found to result in models which best achieved high growth rates while reaching acceptable levels of correlation with input transcriptomes (Table S1).

### Linear Coefficient Assignment

As previously described, remaining reactions were then assigned a linear coefficient based on the user provided transcriptomic abundance data [70]. An option has also been integrated to differentially penalize reactions where multiple gene products are required for a reaction to proceed and discordant transcript abundances are observed based on GPR rules. In order to illustrate this difference, we repeated this analysis with and without GPR consideration (Table S6). The base implementation of RIPTiDe utilizes the maximum transcription abundance level among genes encoding for a single reaction, however the ability to pool overall transcript for all genes contributing to a reaction has also been added.

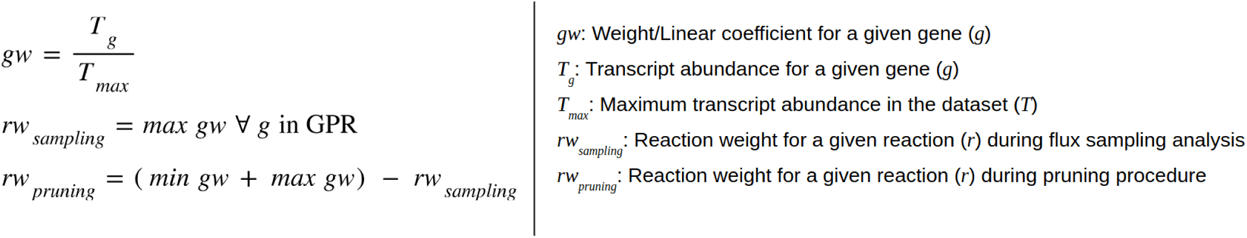

For reaction weight *gw* (linear coefficient) assignments, the reads-per-million normalized transcript abundance of gene *T_g_* within the sorted (smallest to largest) transcript abundance distribution is divided by the largest abundance *T_max_* this distribution. These *gw* values are then assigned to their corresponding reactions to generate a distribution of linear coefficients *rw_sampling_* and the maximum weight among reactions in each gene reaction rule (GPR) is assigned to all associated reactions. The sum of the minimum and maximum weights within *gw* is calculated and then the inverse weight is determined based on the weight associated with reaction *r* in *rw_sampling_*, creating a second distribution of reaction weights *rw_pruning_*.

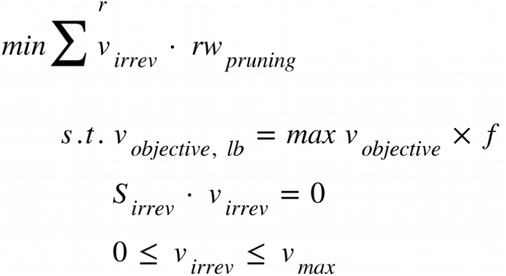

As in parsimonious-FBA, each reversible reaction is separated into individual forward and reverse reactions *v_irrev_*. Linear coefficients *w_r_* for each vector can now be integrated and an objective function is then defined to minimize that total of flux across the entire model. These steps are all performed subject to a lower bound on the biomass flux (*v_objective, lb_*) set to a fraction (*f*, default is 0.8) of the previously identified maximum biomass flux (*max v_objective_*) and mass-balance constraints. Optimization (standard FBA) is performed and reactions found in the solution to have <1x10^-6^ flux are then removed from the model (the threshold is adjustable by the user). The model is then checked for orphaned genes and metabolites, which are subsequently removed as well.

### Flux Sampling

In order to adequately explore intracellular flux through the remaining reactions, previous coefficients are reassigned to remaining reactions using the *rw*_sampling_ linear coefficient assignments. The overall sum of flux is then maximized. The assembly of the large function of weighted reaction fluxes is performed with symengine (https://github.com/symengine/symengine):

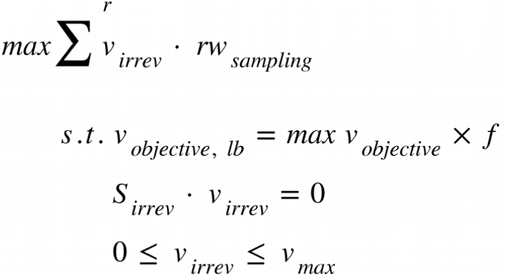

Coefficients (*rw_sampling_*) are then assigned to the corresponding irreversible reactions *v_irrev_*. Flux sampling was then performed with the gapsplit sampler using standard parameters and 500 samples (https://github.com/jensenlab/gapsplit) [41]. Flux variability analysis for the constrained models is also performed to establish flux ranges for each reaction [71]. An overall concordance is then calculated by finding the Spearman correlation coefficient and associated *p*-value between median absolute flux and their associated linear coefficient during flux sampling for all reactions (Table S1 and Fig. S3).

### Example Model Construction

The example model used for initial RIPTiDe testing consists of 16 reactions (7 exchanges, 7 transporters, and 2 metabolic) and 14 metabolites across 2 compartments (extracellular and intracellular). The intracellular demand reaction for ATP generation was utilized as the objective. Simulated transcriptomes are shown in Table S2 and an SBML file (example_GENRE.sbml) for the model is also available at github.com/mjenior/Jenior_RIPTiDe_2019.

### Data Sources and Processing

Fastq files associated with each study were accessed from the NCBI Sequence Read Archive (SRA: PRJNA313438, SRA: PRJNA212627, & SRA: PRJNA415307). Sequencing read processing was performed as described previously in [3, 8]. In summary, raw cDNA reads were quality-trimmed with Sickle [72] at ≧Q30 and subsequently mapped to *Escherichia coli* K-12 MG1655 genes (EMBL-EBI on Jan 28, 2019) using Bowtie2 [73]. MarkDuplicates was then used to remove optical and PCR duplicates (<u>broadinstitute.github.io/picard/</u>). The remaining mappings were converted to idxstats format using SAMtools [74], and the read counts per gene were tabulated. Discordant pair mappings were discarded, and counts were then normalized to read length and gene length to give a per-base report of gene coverage. If replicate sequencing was performed, median transcription values for each gene were then calculated. The data were then converted to a python dictionary of gene keys and associated numeric transcript abundances for use with RIPTiDe. *In vivo* untargeted metabolomic data were generated for a previous study [8], and accessed from <u>github.com/SchlossLab/Jenior_Metatranscriptomics_mSphere_2018/</u>. The current version of iJO1366 (GENRE for *E. coli* K-12 MG1655) was accessed from the BiGG Model database [75] on 09-26-2018.

### Previous Transcriptomic Integration Algorithms

GIMME [11], iMAT [12], MBA [20], CORDA [13], and RegrEx [27] were all implemented using the COBRA toolbox V3 in MATLAB (v9.6; https://opencobra.github.io/cobratoolbox/stable/) using the specified thresholds. For both MBA and RegrEx, a flux consistent model was used based on the requirements for the integration algorithms. CORDA (v0.4.2) and RegRx were downloaded and installed on 12-1-2019.

### Statistical Methods

All statistical analysis was performed using R (v3.2.0). Significant differences between flux distributions and metabolite concentrations were determined by Wilcoxon signed-rank test with Benjamini-Hochberg correction. Principle Component Analysis and Non-Metric Multidimensional Scaling of sampled flux distributions was performed using the vegan package [76]. Machine-learning was accomplished with the implementation of AUC-Random Forest in R [63].

## Data and Software Availability

As previously stated, RIPTiDe is freely available for download at https://github.com/mjenior/riptide and also as a full package (v2.8.0) from the Python Package Index (https://pypi.org/project/riptide/). Complete analysis associated with this study is available in a Jupyter notebook hosted on a GitHub repository along with all other data and code associated with this study (https://github.com/mjenior/Jenior_RIPTiDe_2019).

## Acknowledgements

The authors would like to thank Jhansi Leslie, Greg Medlock, Laura Dunphy, Kristopher Rawls, Maureen Carey, Martin Rau, and Justin Taylor for their suggestions on the utility, execution, and biological validity of the core algorithm. The authors support open-source software development and have constructed this study in line with those principles.

## Supplemental Figures

**Figure S1.**
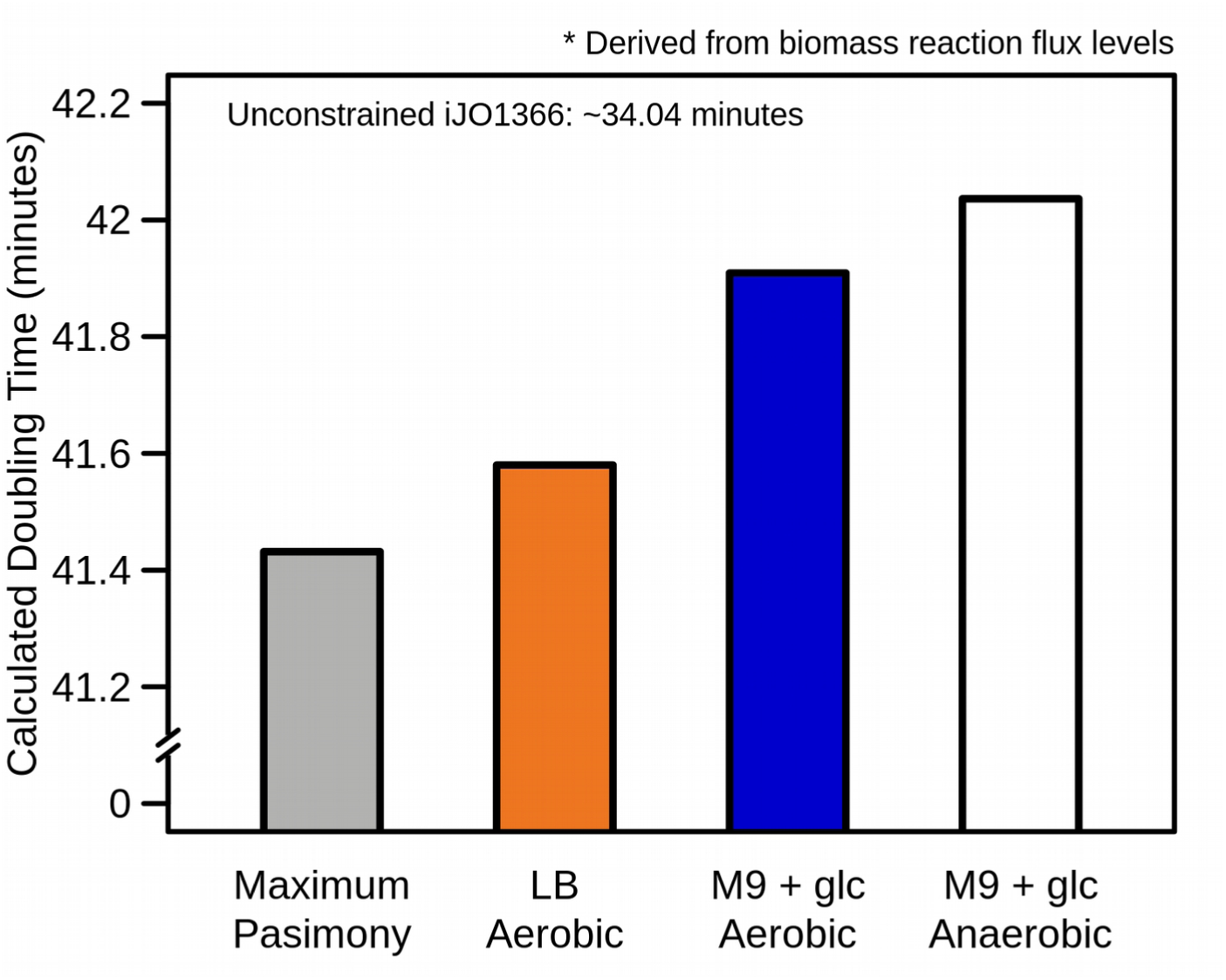
*In silico* growth predictions for *E. coli* K-12 MG1655 supports the utility of RIPTiDe. Computed doubling time from biomass reaction objective value following FBA analysis of each contextualized model (all open exchange reactions). Result from the unconstrained model is listed along the top axis. Biomass objective flux was constrained to ≧80% of the optimal flux value prior to identifying the state of Maximum Parsimony as it is during RIPTiDe contextualization.

**Figure S2.**
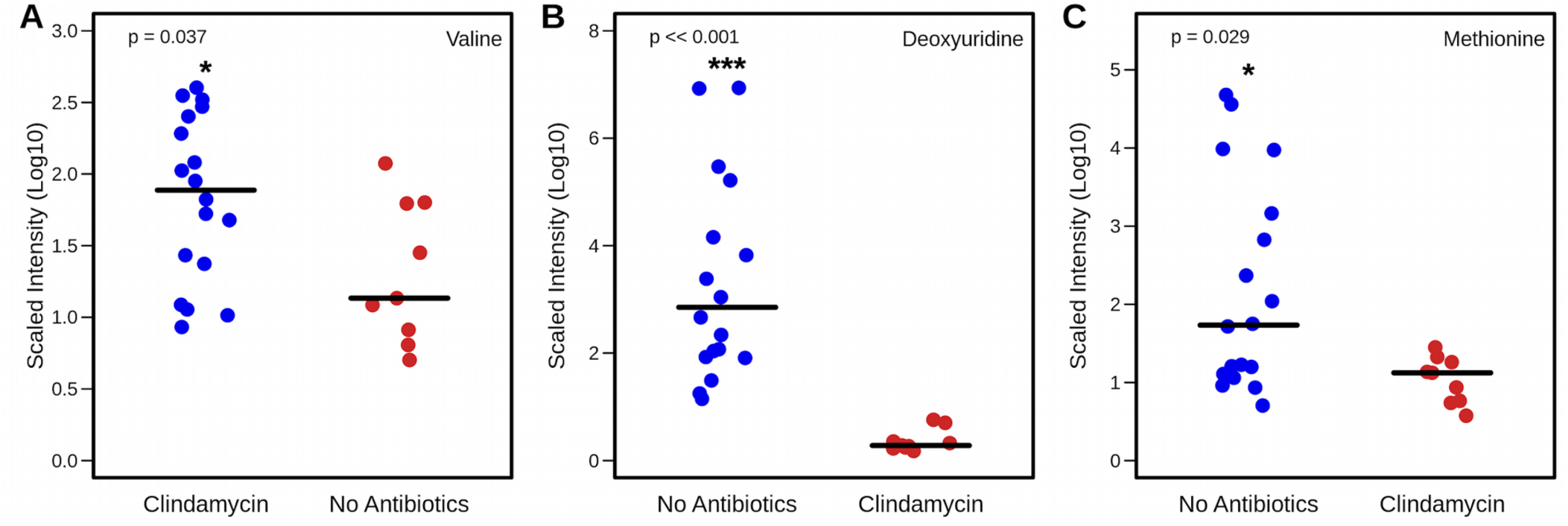
Relative concentrations of metabolites highlighted during *in vivo* analysis of iJO1366 with RIPTiDe. Concentrations quantified with liquid chromatography mass-spectrometry previously [3]. Comparing the metabolite content in cecal content of mice pretreated with an intraperitoneal injection of clindamycin against untreated control animals. **(A)** Valine, **(B)** Deoxyuridine, **(C)** and Methionine. Significant differences were determined by Wilcoxon signed-rank test.

**Figure S3.**
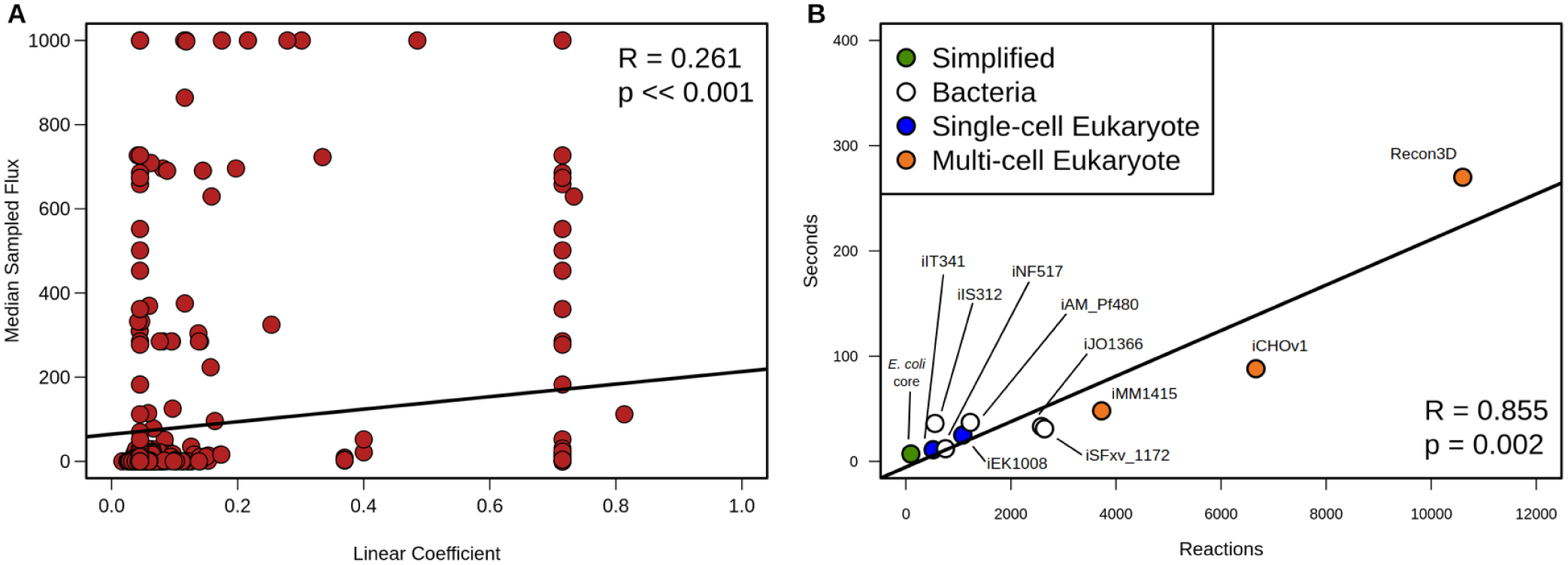
RIPTiDe performance metrics. **(A)** Spearman correlation results between RIPTiDe linear coefficients from flux sampling and median absolute values from sampled flux ranges from *in vivo* transcriptome. Assigned linear coefficients are significantly correlated with their associated absolute reaction activities. Correlation results are referred to as concordance in the output of RIPTiDe. **(B)** Run time in seconds with increasing metabolic network reconstruction size. Using RIPTiDe, each GENRE (11 total) was integrated with a simulated transcriptome of 1.0 transcript for all genes. Multicellular eukaryotic models include mouse (iMM1415), golden hamster (iCHOv1), and human (Recon3D). The relationship between model size and RIPTiDe run time fits a linear model (*p*-value = 0.002). These analyses were completed using an Intel® Core™ i7-7Y75 CPU @ 1.30GHz × 4 processor and 15.4 GB of memory.

**Figure S4.**
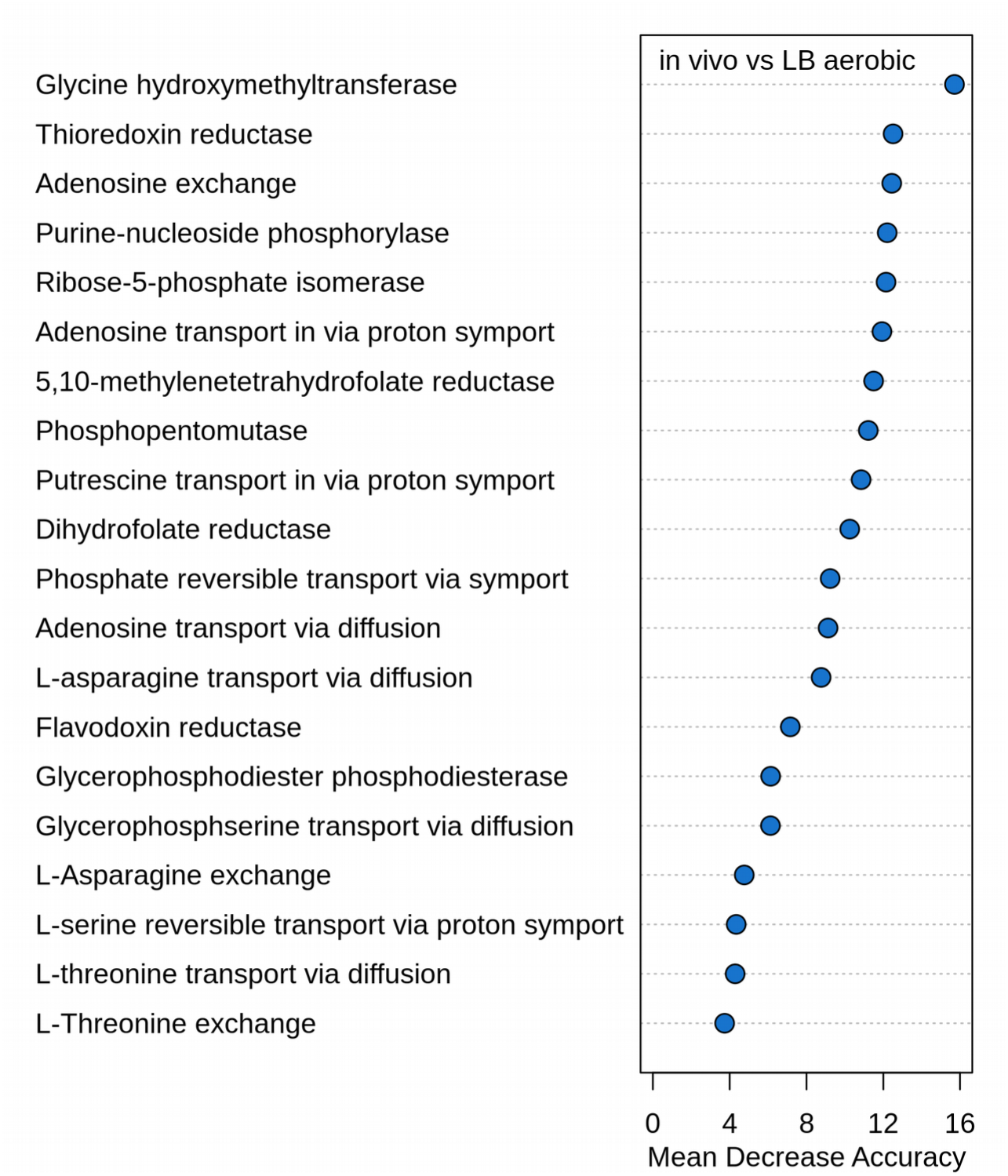
Results from Random Forest machine learning to identify the most differentiating metabolic reactions between models. Mean Decrease Accuracy for an optimal subset of 20 reactions determined by AUC Random Forest that flux levels differentiate the distributions from shared reactions between *in vivo* and LB rich media RIPTiDe-contextualized transcriptomes of *E. coli*.

**Figure S5.**
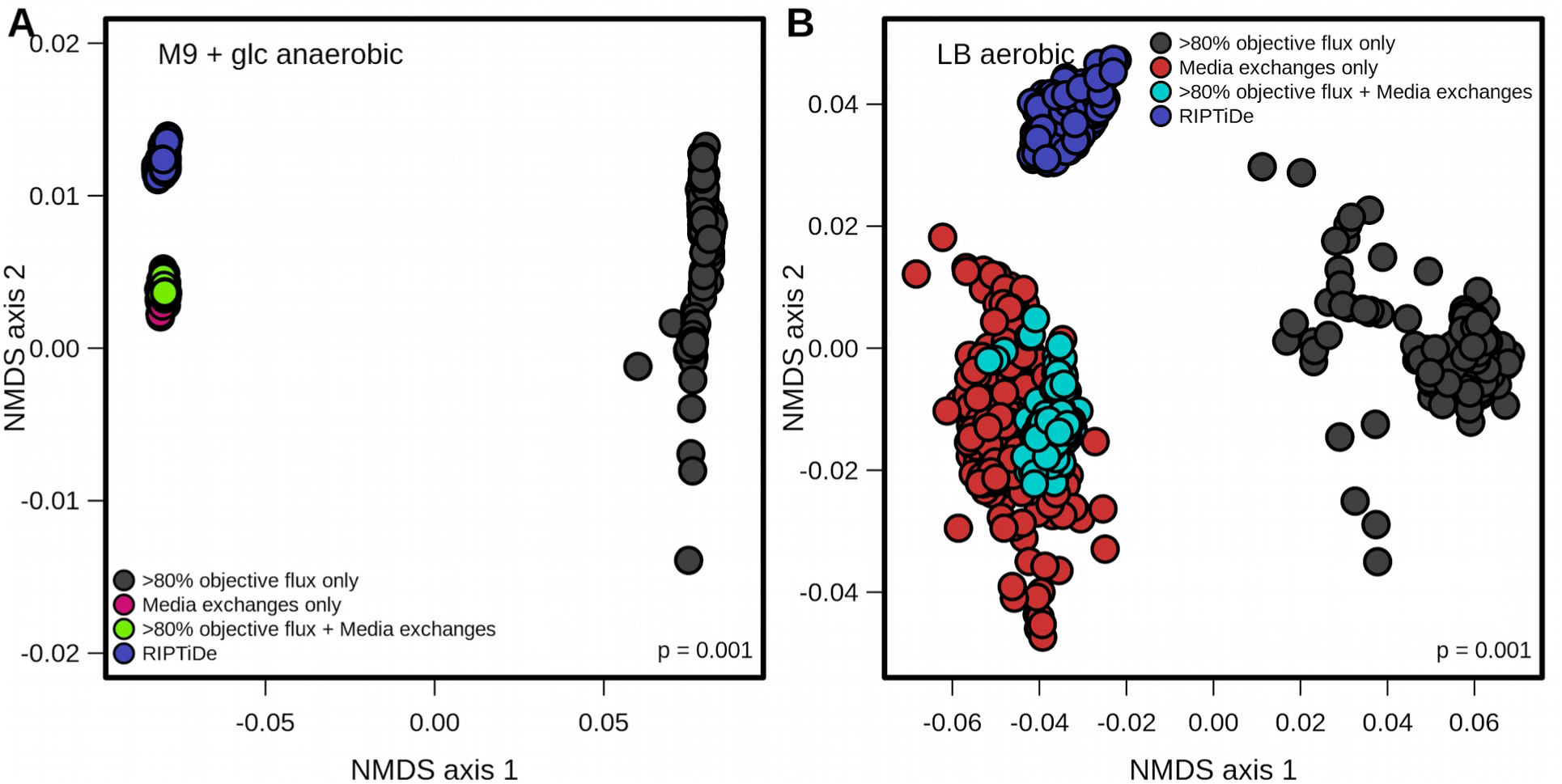
Flux samples from context-specific models with RIPTiDe vs exchange flux media constraints follow the same trends across all conditions tested. NMDS ordination of Bray-Curtis dissimilarities between flux samples for each version of iJO1366 in (A) M9+glucose anaerobic and (B) LB aerobic. Significant differences between constrained objective flux only and all other groups determined by PERMANOVA. The dissimilarity of RIPTiDe-generated flux samples in these analyses highlight context-specific metabolic patterns that were unobservable through other methods.

Table S1 | Sensitivity testing for minimum objective threshold percentage

Table S2 | Structure and analysis for example metabolic reconstruction

Table S3 | Parameters and results from previous transcriptomic integration algorithms

Table S4 | Bray-Curtis dissimilarities between flux distributions from differentially constrained versions of iJO1366

Table S5 | Network topology and gene essentiality across RIPTiDe contextualized models

Table S6 | Comparison of RIPTiDe Models Resulting from LB-media Transcriptome Contextualization With and Without Consideration for Gene-Protein Reaction Rules

Table S7 | Growth Substrate Predictions Across Transcriptome Integration Algorithms

